# The Consequences of Egg Adaptation in the H3N2 Component to the Immunogenicity of Live Attenuated Influenza Vaccine

**DOI:** 10.1101/834887

**Authors:** Daniel H. Goldhill, Benjamin Lindsey, Ruthiran Kugathasan, Zandra Felix Garza, Ya Jankey Jagne, Hadijatou Jane Sallah, Gabriel Goderski, Sophie van Tol, Katja Höschler, Adam Meijer, Wendy S. Barclay, Thushan I. de Silva

## Abstract

Adaptation in egg-passaged vaccine strains may cause reduced vaccine effectiveness due to altered antigenicity of the influenza haemagglutinin. We tested whether egg adaptation modified serum and mucosal antibody responses to the A(H3N2) component in the Live Attenuated Influenza Vaccine (LAIV). Twice as many children seroconverted to an egg-adapted H3N2 than the equivalent wildtype strain. Seroconversion to the wildtype strain was greater in children seronegative pre-LAIV, whereas higher mucosal IgA responses to wildtype antigen were observed if seropositive prior to vaccination. Sequencing of virus from nasopharyngeal swabs from 7 days post-LAIV showed low sequence diversity and no reversion of egg-adaptive mutations.

## Background

Vaccines offer the best protection against morbidity and mortality caused by influenza. Low vaccine effectiveness (VE) is often attributed to an antigenic mismatch between the vaccine and circulating strains caused by evolution of the influenza haemagglutinin (HA) [1]. Recently, despite a well-matched vaccine strain selection, low VE of the A(H3N2) component was reported, potentially due to adaptive mutations caused by the production of the influenza vaccine in eggs [1]. These egg-adaptive mutations can alter vaccine antigenicity and lead to immune responses mismatched to circulating strains.

Of the four components of the current quadrivalent seasonal influenza vaccine, the issue of egg adaptation resulting in antigenic mismatch is most problematic for H3N2 subtype influenza A viruses. In 2014, the H3N2 3C.2a clade arose with a K160T mutation, giving an additional putative glycosylation site in immunodominant antigenic site B on the HA head [2]. In order to grow efficiently, 3C.2a strains revert from T160 to K160 when cultured in eggs and this adaptive mutation has been suggested as a key reason for the recent loss of H3N2 VE [1]. The reduction in immunogenicity against circulating strains caused by this reversion has so far been described in the context of serum antibody responses to inactivated influenza vaccine (IIV) [1]. Here, we examine the consequence of egg-adaptations in the Live Attenuated Influenza Vaccine (LAIV) on serum and mucosal antibody responses in young children to circulating T160-containing H3N2. We also sequence virus recovered from the nasopharynx after LAIV immunisation to test whether there is reversion to a human-adaptive form.

## Methods

### Study design and sample collection

The samples used in this study were generated during a larger randomised controlled trial (ClinicalTrials.gov NCT02972957) comparing the impact of LAIV on the nasopharyngeal microbiome. The trial was conducted in Sukuta, a periurban area in The Gambia during 2017 and 2018. A detailed description of the cohort and sampling is described elsewhere [3]. Influenza vaccine-naïve children aged 24-59 months received a single intranasal dose of the trivalent Russian-backbone LAIV (Nasovac-S, Serum Institute of India Pvt Ltd) containing the World Health Organisation recommended viruses for the Northern Hemisphere for either 2016-2017 or 2017-2018, dependent on the year of enrolment. For both vaccines, the H3N2 component was an A/Hong Kong/4801/2014 (H3N2)-like virus. H3N2 vaccine titres per dose (50% Egg Infectious Doses (EID50)/ml) were 1×10^7.5^ in 2017 and 1×10^7.6^ in 2018. Nasopharyngeal swabs (FLOQSwabs, Copan, Murrieta, CA, USA) were collected into RNAprotect cell reagent (Qiagen) on day 2 (D2) and 7 (D7) post-vaccination. Serum and oral fluid swab (Oracol Plus; Malvern Medical Development, Worcester, UK) samples were taken on D0 and day 21 (D21). All samples were stored at −70°C before further processing. The study was approved by The Gambia Government/MRC Joint Ethics Committee and the Medicines Control Agency of The Gambia. A parent provided written or thumb-printed informed consent for their children to participate.

### HAI and IgA assays

Haemagglutinin inhibition (HAI) assays were carried out on serum samples using guinea pig red blood cells (0.5% in PBS) in the absence of oseltamivir and with standard methods [4]. HAI titres pre- and D21 post-LAIV were determined to cell-cultured (in MDCK-SIAT cells) and egg-cultured A/Hong Kong/4801/2014 (HK14) strains. Sequencing of viral stocks confirmed the presence of T160 in cell-cultured and K160 in egg-cultured HK14. Egg-cultured virus also contained further egg-adaptive mutations L194P, T203I and I260L. Seroconversion was defined as a 4-fold rise in HAI titre to ≥1:40, irrespective of baseline HAI titre, using a 2-fold dilution series of serum starting from 1:10 dilution. Mucosal influenza-specific IgA was measured in oral fluid samples at baseline and D21 post-LAIV using a protein microarray as previously described [5, 6] with the percentage Surfact-Amps-20 in the blocking, washing and incubation buffer increased from 0.05% to 5% to prevent background staining with oral specimens. Microarrays were coated with recombinant HA1 protein expressed in human cells (Sino Biological, Beijing, China) reflecting sequences of egg-adapted and cell-cultured HK14. Total IgA was quantified using an ELISA and influenza-specific IgA expressed as a ratio of influenza HA1-specific IgA/total IgA as previously described [6].

### RNA Extraction, Primer ID and Sequencing

RNA was extracted from D2 and D7 nasopharyngeal samples previously identified as positive for H3N2 shedding by RT-PCR [3], using QIAamp Viral RNA Mini Kit (Qiagen) with carrier RNA. RNA was also extracted from vaccine aliquots diluted 1000-fold. RNA was reverse transcribed using Superscript III (Invitrogen) and a barcoded primer specific to each 500bp sub-amplicon in the HA (Primer ID). Primer ID attaches a unique barcode to each cDNA molecule during reverse transcription and allows for PCR and sequencing error correction [7, 8]. PCR was performed using KOD polymerase (Merck). Samples were pooled across sub-amplicons and prepared for sequencing using NebNext Ultra II (NEB), then sequenced on an Illumina MiSeq with 300bp paired-end reads. Sequences were analysed in Geneious (v11) and a pipeline in R. Forward and reverse reads were paired using FLASh (https://ccb.jhu.edu/software/FLASH) before being mapped to a reference sequence and consensus sequences made for each barcode. Degenerate barcodes were removed (see Supplementary material and Figure S1) and a minimum cut-off of 5 reads per barcode was chosen. Raw sequences were deposited at https://www.ebi.ac.uk/ena (project number PRJEB34129.) The analysis pipeline can be found at https://www.github.com/Flu1/GambiaLAIV.

### Statistical analysis

Paired and unpaired proportions were compared using McNemar’s and Chi^2^ tests respectively. HAI geometric mean fold rise (GMFR) within and between individuals was compared using the Wilcoxon signed-rank and Mann-Whitney tests. Log10-transformed IgA fluorescence ratio fold-change was compared using paired and unpaired t-tests. Pairwise correlations were assessed using Spearman’s rank-order (GMFR) and Pearson’s (log10-transformed IgA fold-change) test. Shannon Entropy was used to calculate the diversity of mutations within each sequenced sample (Supplementary material). Genetic distance between samples was calculated as described in Supplementary material. Statistical analyses were performed using R version 3.5.0 and GraphPad Prism 8.0.2.

## Results

Samples from 244 children were included in the HAI analysis (n=118 from 2017 and n=126 from 2018 [3]). Influenza-specific oral fluid IgA data were available from 214 children (n=100 from 2017 and n=114 from 2018). D7 nasopharyngeal swab samples from 30 children with H3N2 detected by RT-PCR along with D2 samples from 22/30 were available for sequencing.

No significant differences were seen between pre-LAIV HAI titres to egg-cultured and cell-cultured HK14 (p=0.84, Figure S2). The proportion of children who seroconverted to egg-cultured HK14 virus was 25.0% (61/244, 95% confidence interval (CI) 19.6-30.4) compared to 12.3% (30/244, 95% CI 8.2-16.4) to cell-cultured HK14 (p<0.0001). D0 to D21 GMFR to egg-cultured HK14 was greater than to cell-cultured HK14 (p<0.0001, Figure 1A). A significant correlation was present between GMFR to egg-cultured and cell-cultured HK14 (r_s_=0.58, p<0.0001, Figure S3), although discrepant samples were observed with seroconversion to only one virus (Figure 1B). In contrast, the increase in mucosal influenza HA-specific IgA from D0 to D21 post-LAIV was greater to cell-culture matched HK14 HA compared to egg-culture matched HK14 HA (p=0.0009, Figure 1C). A significant correlation was observed between IgA fold-change to egg- and cell-cultured HK14 HA (r=0.69, p<0.0001, Figure S4).

**Figure 1.**
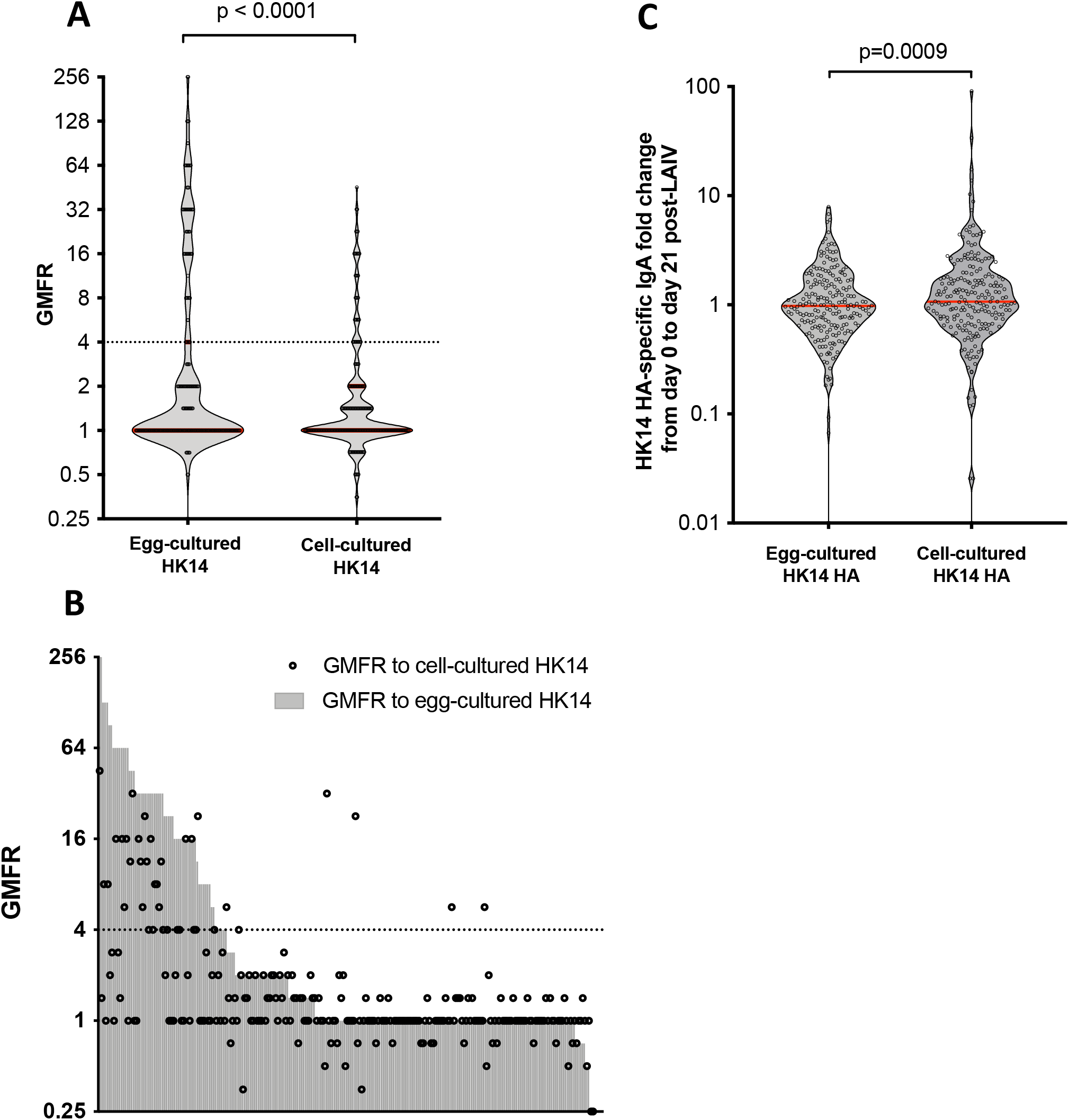
**a)** Geometric Mean Fold Rise (GMFR) to egg-cultured and cell-cultured HK14 **b)** Fold-change from day 0 to day 21 post-LAIV in HA1-specific IgA to proteins representing egg-cultured and cell-cultured HK14 **c)** GMFR to egg and cell-cultured HK14 for each individual. Dotted line represents 4-fold increase in HAI titre between day 0 and day 21 which defines seroconversion.

To explore the impact of prior H3N2 infection on serum and mucosal antibody responses to HK14 in LAIV, children were stratified based on seropositivity to cell-cultured HK14 (pre-LAIV HAI titre ≥1:10). In seronegative children, seroconversion to egg-cultured HK14 (50.7%, 37/73, 95% CI 39.2-62.2%) and cell-cultured HK14 (27.4%, 20/73, 95% CI 17.2-37.6%) was greater than in seropositive children (14.0% seroconversion to egg-cultured, 24/171, 95% CI 8.8-19.4%, p=0.00048 and 5.8% seroconversion to cell-cultured, 10/171, 95% CI 2.3-9.4%, p<0.0001). This pattern was reflected in GMFR values (Figure 2A), but not in IgA responses. D0 to D21 post-LAIV IgA fold change, to HK14 HA proteins representative of both egg-cultured and cell-cultured HK14, was modestly higher in seropositive children compared to seronegative children (Figure 2A).

**Figure 2.**
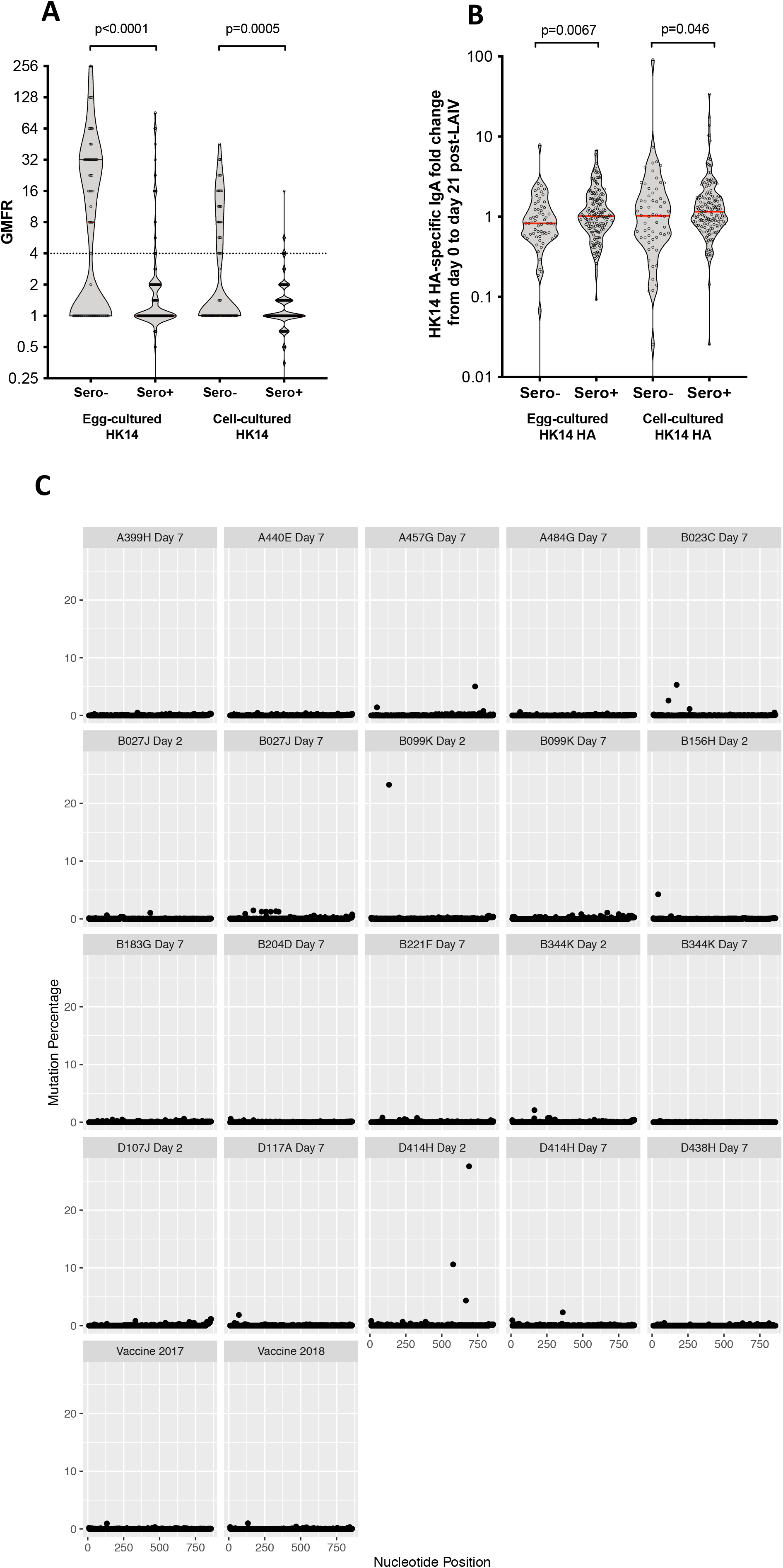
**a)** GMFR to egg and cell-cultured HK14 comparing children seropositive and seronegative to cell-cultured (i.e. wild-type) HK14 prior to receiving LAIV. Dotted line represents 4-fold increase in HAI titre between day 0 and day 21 which defines seroconversion. **b)** Fold-change from day 0 to day 21 post-LAIV in HA1-specific IgA to proteins representing egg-cultured and cell-cultured HK14, comparing seropositive and seronegative children. **c)** Shed virus from 20 samples from either day 2 or day 7 and the vaccine from 2017 and 2018 were sequenced using Primer ID. The percentage of mutations are shown at each sequenced nucleotide position in the HA where 1 refers to the first base of the signal peptide. The sample ID and day of sample collection are shown.

To explore whether reversion of egg-adapted mutations in LAIV HK14 during nasopharyngeal replication could drive responses to cell-cultured HK14, we sequenced two sub-amplicons containing HA amino-acids 1-276. 20 samples with low H3N2 cycle threshold (Ct) values (five D2 and fifteen D7 from sixteen individuals) were successfully amplified (Supplementary material, Figure S5). No significant reversion of egg-adaptive mutations was seen in any samples (Figure 2C). This included D7 samples from seven seroconverters to cell-cultured HK14. Two samples showed a single sequence with P194L (<0.2% frequency) and three samples showed one or two sequences with I203T (<0.2% frequency). Along with position 160, these three sites were >99.9% identical to the vaccine across all samples. Few mutations rose to high frequency with only five mutations occurring above 5%. Of these, I23L was a pre-existing polymorphism present at 1% in the vaccine strain and the other four mutations were synonymous. There was no significant difference between Shannon entropy for the samples and vaccine strains (Z-test, p=0.41). In individuals with matched samples, mutations present at higher frequencies on D2 had been lost by D7 (Figure 2C). Using the relative L1-norm as a measure of genetic similarity, there was no significant difference between samples taken from the same individual and other samples (Z-test, p=0.54, Figure S6).

## Discussion

We describe, to our knowledge for the first time, the impact of egg-adaptations in a recent H3N2 3C.2a strain vaccine antigen on serum and mucosal antibody responses induced by LAIV to the equivalent human-adapted strain reflective of circulating viruses. In keeping with observations with IIV [2], serum antibody responses to cell-cultured HK14 were significantly lower than to the vaccine-matched egg-cultured HK14. However, a proportion of children did seroconvert to cell-cultured HK14, which was most evident in children seronegative to cell-cultured HK14 prior to receiving LAIV. In the absence of prior HK14 exposure, the serum antibody response in these children may be broader and directed to antigens outside antigenic site B [2].

In contrast to serum HAI induction, IgA responses to proteins representing cell-cultured HK14 HA were equivalent or higher than those representing egg-cultured HK14 HA. IgA responses were also modestly higher in children who were seropositive to HK14 prior to LAIV compared to seronegative children. Therefore, in our cohort, unlike serum antibodies induced by LAIV, mucosal IgA responses may largely reflect boosting of prior responses acquired through natural infection. However, the lack of a significant IgA correlate of protection following LAIV vaccination means that the clinical relevance of this finding is uncertain. Compared to the serum HAI response, little is known about the antigen-specificity of LAIV-induced mucosal IgA responses, although some studies have suggested influenza-specific IgA responses are more cross-reactive that IgG responses [9, 10]. It is important to note that the IgA responses measured constitute binding antibody, rather than functional responses, which are challenging to measure in mucosal samples. Future work could explore functional mucosal IgA responses, as well as anti-HA stalk responses which we did not assess and may provide cross-reactive responses.

A previous influenza human challenge study in adults has demonstrated the reversion of an egg-adapted mutation during replication in the upper respiratory tract [11]. We hypothesized that a similar phenomenon could occur with LAIV replication of HK14, providing a potential explanation for cell-cultured HK14-specific antibody responses after vaccination with an egg-adapted antigen. Sequencing of the shed virus, however, revealed no changes at sites of egg adaptation and very few significant changes in the HA. The lack of a K160 fitness cost in humans is perhaps unsurprising given the majority of H3N2 isolates prior to 2014 contained K160. Recent studies have found low within-host diversity of virus in natural influenza infections in vaccinated and unvaccinated individuals, suggesting that the immune system does not put significant pressure on the influenza virus to evolve over the course of an individual infection [12, 13]. Our results agree with this and imply there is little positive selection on the LAIV H3N2 HA in the nasopharynx within the first week and that reversion of egg-adaptation mutations such as K160 is unlikely.

Although egg adaptation is likely to be an important factor, increasing data suggest several factors contribute to the low VE to H3N2 observed in some years [14]. Developing alternatives to egg-based methods of vaccine production is clearly important as current vaccines may result in protective H3N2 responses only in sub-populations of individuals.

## Funding

This work was supported by a Wellcome Trust Intermediate Clinical Fellowship award to TdS (110058/Z/15/Z) and by a Biotechnology and Biological Sciences Research Council grant (BB/K002465/1) and a Wellcome grant (205100/Z/16/Z) to WSB. RK was supported by Wellcome fellowship (216353/Z/19/Z).

## Acknowledgements

We gratefully acknowledge the study participants and parents who took part in the study. We also acknowledge the dedicated field team involved in carrying out the clinical trial. We thank Christine Carr, Monali Patel and Hirushi Rajapakse for their technical support at the Respiratory Virus Unit (PHE NIS, Colindale).

## Notes

**Conflicts of Interest:** The authors have no conflicts of interest.

