## Supplementary Data for "The Consequences of Egg Adaptation in the H3N2 Component to the Immunogenicity of Live Attenuated Influenza Vaccine"

### Supplementary material

#### Degenerate Barcodes

Sequencing errors in the barcode region can lead to novel, degenerate barcodes. Some samples contained high numbers of reads for individual barcodes which increases the likelihood of degenerate barcodes with reads greater than the cut-off for minimum number of reads. A published formula for removing degenerate barcodes sets a threshold determined by the barcode with the highest number of reads[1]. However, we found this formula was inaccurate when samples contain a barcode with very high read numbers (>10,000 reads) and imposing a strict cut-off resulted in the loss of many sequences which do not appear degenerate[1]. Therefore, we wanted a more precise method for removing degenerate barcodes. We compared the distribution of barcode sequences to a random distribution of barcode sequences and found a 165-fold overrepresentation of barcodes separated by 1 mutation and a 23-fold overrepresentation of barcodes separated by 2 mutations (Fig S1A). Comparing the respective number of reads for barcodes separated by 1 or 2 mutations confirmed that this overrepresentation was caused by large numbers of degenerate barcodes with a low number of reads 1 or 2 mutations away from a barcode with a high number of reads (Fig S1B). Therefore, we removed all such degenerate barcodes from the analysis.

#### Shannon Entropy and genetic distance between samples

Shannon Entropy was used to calculate the diversity of mutations within a sample according to the following formula: $H(x) = \sum_{i}^{s} P_i log_2 P_i$ across *s* total sites where $P_i$ is the relative frequency of a variant at position *i*. This was implemented using Entropy from the DescTools

(v. 0.99.28) package in R. To calculate genetic distance between samples, we used the L1-norm $L = \sum_i^s \sum_j^n |P_{i,j} - Q_{i,j}|$, where $P_{i,j}$ and $Q_{i,j}$ are the relative frequency of the nucleotide *j* at position *i* summed across *n* nucleotides (A,C,G,T,-) and *s* sites for samples *P* and *Q* respectively. To adjust for the difference in number of mutations within each sample, we normalized across all the samples by dividing each sample by the average of the L1-norm between that sample and all other samples.


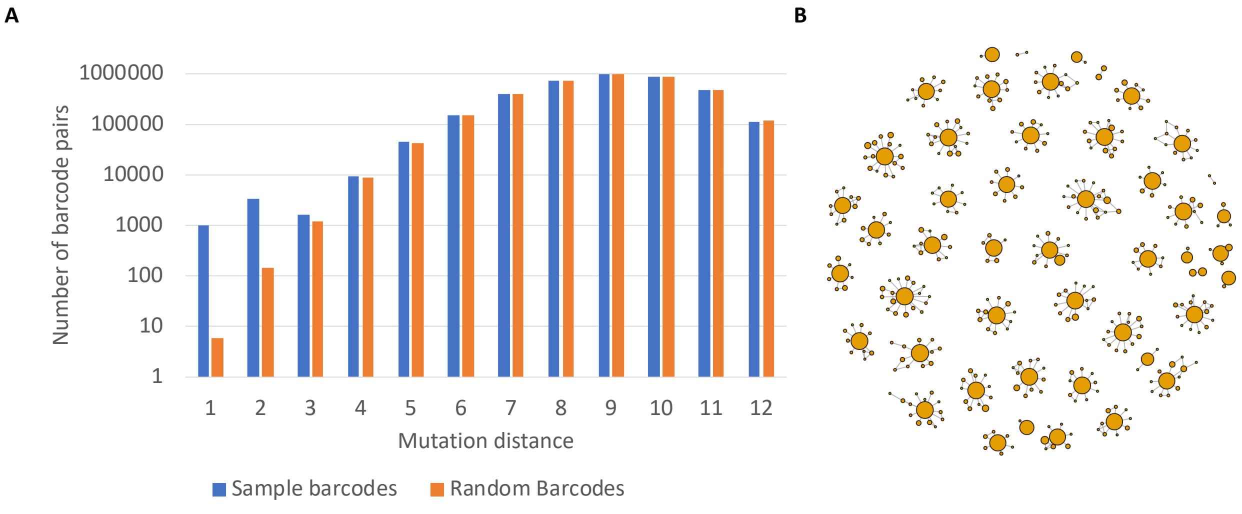


**Fig S1. Degenerate barcodes are produced by barcodes with high numbers of reads. A)** The number of mutations between all pairs of barcodes was calculated for each sample and for an equivalent number of randomly generated barcodes. The number of barcode pairs was plotted against the number of mutations separating the barcode pair. **B)** Barcode pairs were plotted which were separated by a single mutation. The size of circle is the log of the number of reads per barcode. There were 988 barcode pairs in the sample displayed compared to 6 expected barcode pairs from the random sample.

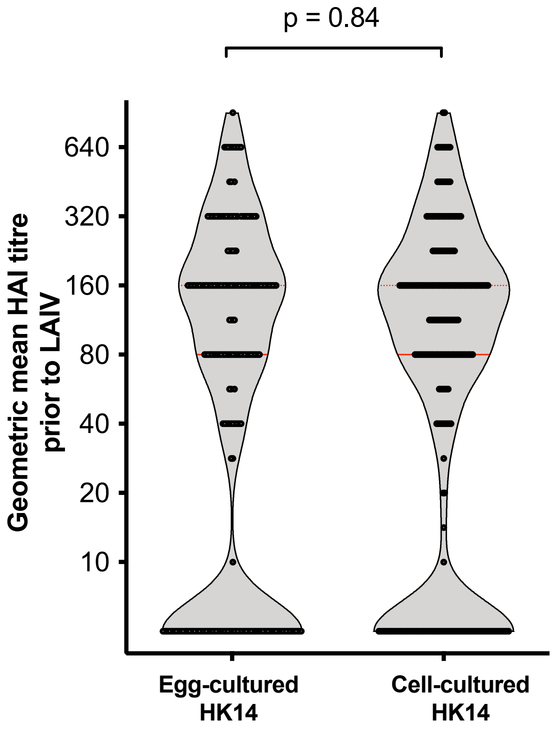


Fig S2. Haemagglutination inhibition (HAI) titres to egg-cultured HK14 and cell-cultured HK14 in serum samples taken prior to LAIV.

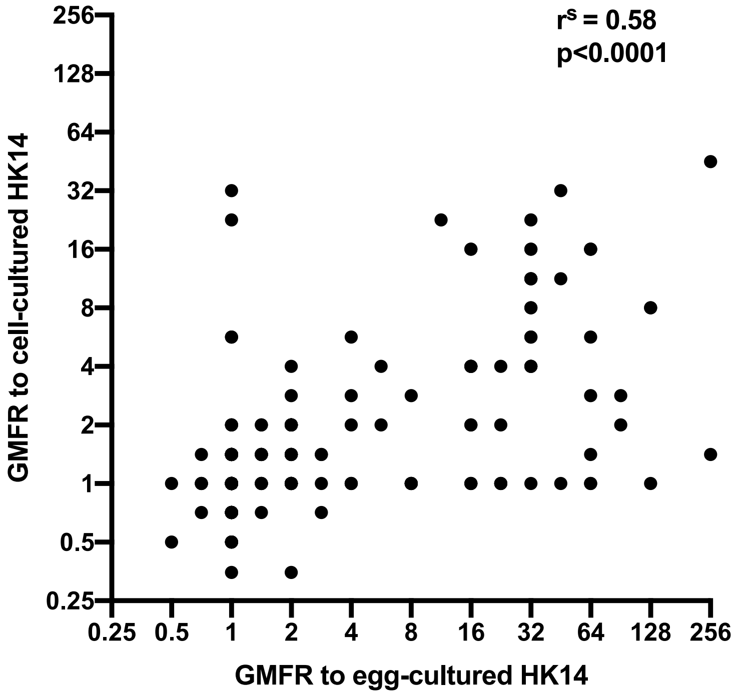


Fig S3. Correlation between geometric mean fold rise (GMFR) in haemagglutination inhibition titre to H3N2 cell-cultured HK14 and egg-cultured HK14 A(H3N2) viruses.

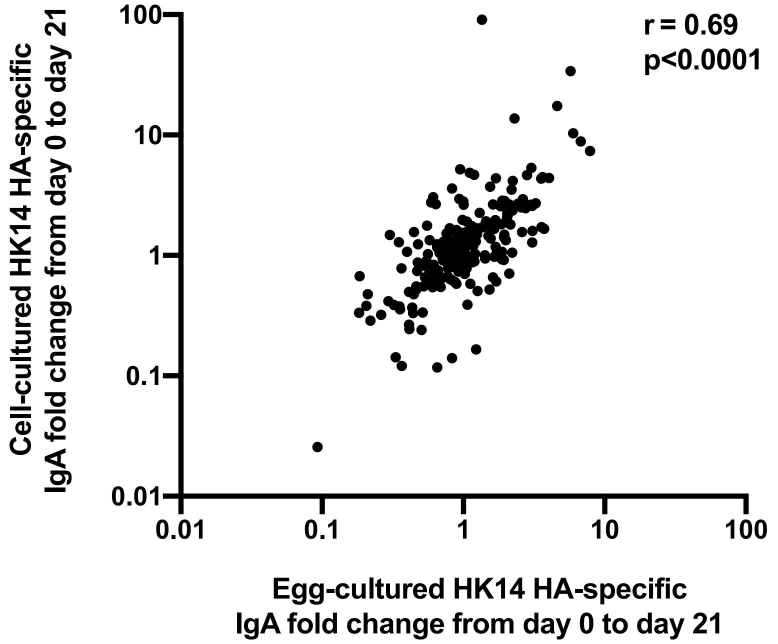


Fig S4. Correlation between egg-cultured HK14 HA-specific and cell-cultured HK14 HA1-specific IgA fold change between day 0 and day 21 post-LAIV. Influenza-specific IgA was normalised by total IgA at each timepoint prior to calculation of fold change. HA1 = globular head of haemagglutinin.

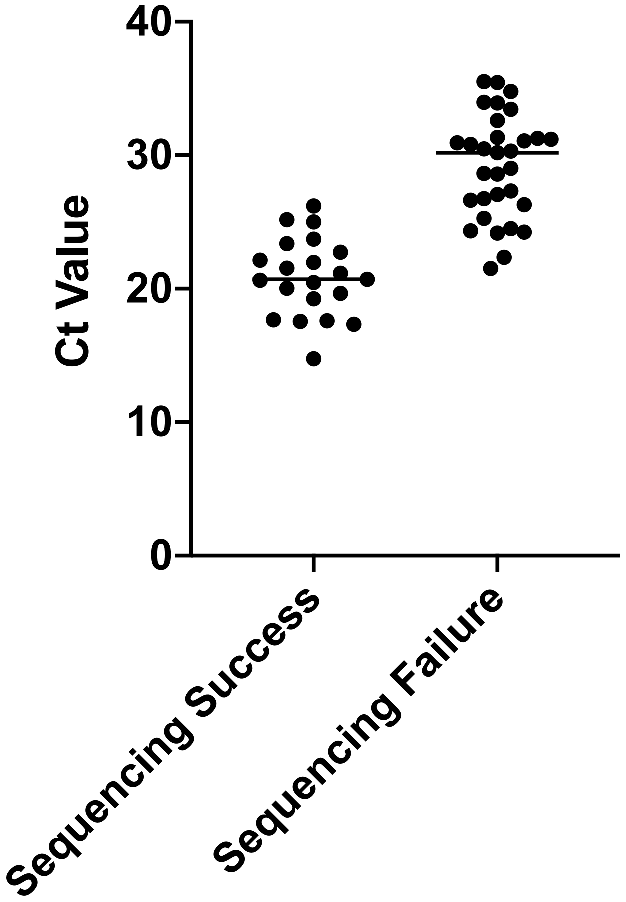


**Fig S5.** Ct values for the M segment from nasal samples were previously measured. This figure shows whether HA subamplicons were successfully amplified and sequenced. On average 2007 barcoded molecules were successfully sequenced for each sub-amplicon for each sample (min = 322, max = 5,104, s.d. = 1,342). One sample with <10 barcodes excluded from the analysis.

| Relative L1 Norm | B027J Day 2 | B099K Day 2 | B156H Day 2 | B344K Day 2 | D107J Day 2 | D414H Day 2 | A399H Day 7 | A440E Day 7 | A457G Day 7 | A484G Day 7 | B023C Day 7 | B027J Day 7 | B099K Day 7 | B183G Day 7 | B204D Day 7 | B221F Day 7 | B344K Day 7 | D117A Day 7 | D414H Day 7 | D438H Day 7 | Vaccine 2017 | Vaccine 2018 |
|---|---|---|---|---|---|---|---|---|---|---|---|---|---|---|---|---|---|---|---|---|---|---|
| **B027J Day 2** | | 1.04 | 0.92 | 1.03 | 1.06 | 1.01 | 1.00 | 0.98 | 0.93 | 0.91 | 0.93 | 1.01 | 1.06 | 0.96 | 0.96 | 0.89 | 0.84 | 0.88 | 0.89 | 0.90 | 0.82 | 0.85 |
| **B099K Day 2** | 1.94 | | 1.89 | 1.67 | 1.40 | 1.43 | 1.91 | 1.86 | 1.91 | 2.18 | 1.90 | 1.66 | 1.44 | 2.00 | 2.17 | 2.10 | 2.38 | 2.08 | 2.06 | 2.23 | 1.96 | 1.99 |
| **B156H Day 2** | 0.91 | 1.00 | | 0.96 | 0.97 | 0.99 | 0.93 | 0.89 | 0.98 | 0.89 | 0.94 | 1.01 | 0.97 | 0.86 | 0.81 | 0.93 | 0.91 | 0.94 | 0.94 | 0.92 | 0.91 | 0.90 |
| **B344K Day 2** | 1.08 | 0.95 | 1.03 | | 0.89 | 1.03 | 0.88 | 1.01 | 1.08 | 1.05 | 1.06 | 0.92 | 0.93 | 1.01 | 1.02 | 1.04 | 1.07 | 1.07 | 1.06 | 1.07 | 1.06 | 1.07 |
| **D107J Day 2** | 1.32 | 0.94 | 1.22 | 1.05 | | 1.13 | 1.15 | 1.20 | 1.27 | 1.34 | 1.28 | 1.15 | 0.91 | 1.16 | 1.28 | 1.34 | 1.41 | 1.33 | 1.35 | 1.36 | 1.35 | 1.33 |
| **D414H Day 2** | 2.23 | 1.70 | 2.22 | 2.16 | 2.00 | | 2.35 | 2.28 | 2.12 | 2.55 | 2.18 | 2.01 | 2.14 | 2.50 | 2.61 | 2.45 | 2.74 | 2.38 | 2.33 | 2.56 | 2.33 | 2.37 |
| **A399H Day 7** | 0.88 | 0.91 | 0.84 | 0.74 | 0.82 | 0.94 | | 0.79 | 0.88 | 0.80 | 0.86 | 0.78 | 0.79 | 0.78 | 0.77 | 0.82 | 0.81 | 0.84 | 0.83 | 0.84 | 0.89 | 0.88 |
| **A440E Day 7** | 0.92 | 0.94 | 0.85 | 0.89 | 0.91 | 0.96 | 0.83 | | 0.87 | 0.85 | 0.90 | 0.94 | 0.85 | 0.78 | 0.76 | 0.87 | 0.90 | 0.93 | 0.90 | 0.92 | 0.92 | 0.89 |
| **A457G Day 7** | 1.01 | 1.12 | 1.09 | 1.12 | 1.12 | 1.05 | 1.09 | 1.01 | | 1.03 | 1.05 | 1.09 | 1.12 | 1.12 | 1.09 | 1.02 | 0.89 | 0.96 | 1.04 | 0.91 | 0.99 | 1.00 |
| **A484G Day 7** | 0.72 | 0.93 | 0.72 | 0.79 | 0.86 | 0.91 | 0.72 | 0.72 | 0.74 | | 0.71 | 0.82 | 0.82 | 0.66 | 0.59 | 0.64 | 0.55 | 0.68 | 0.65 | 0.61 | 0.67 | 0.69 |
| **B023C Day 7** | 0.97 | 1.07 | 1.00 | 1.05 | 1.08 | 1.04 | 1.02 | 1.01 | 1.01 | 0.95 | | 0.88 | 1.07 | 1.03 | 0.97 | 0.97 | 0.98 | 1.00 | 0.96 | 1.00 | 0.98 | 1.02 |
| **B027J Day 7** | 1.22 | 1.09 | 1.24 | 1.06 | 1.12 | 1.10 | 1.07 | 1.22 | 1.21 | 1.26 | 1.01 | | 1.11 | 1.24 | 1.28 | 1.18 | 1.33 | 1.26 | 1.21 | 1.30 | 1.23 | 1.24 |
| **B099K Day 7** | 1.16 | 0.85 | 1.08 | 0.97 | 0.80 | 1.06 | 0.98 | 1.00 | 1.12 | 1.13 | 1.12 | 1.00 | | 0.97 | 1.07 | 1.15 | 1.19 | 1.17 | 1.17 | 1.17 | 1.19 | 1.18 |
| **B183G Day 7** | 0.78 | 0.87 | 0.71 | 0.78 | 0.76 | 0.92 | 0.72 | 0.67 | 0.83 | 0.68 | 0.79 | 0.83 | 0.72 | | 0.56 | 0.72 | 0.69 | 0.74 | 0.73 | 0.69 | 0.80 | 0.77 |
| **B204D Day 7** | 0.72 | 0.87 | 0.62 | 0.73 | 0.77 | 0.88 | 0.65 | 0.60 | 0.74 | 0.55 | 0.69 | 0.79 | 0.73 | 0.52 | | 0.63 | 0.54 | 0.65 | 0.63 | 0.58 | 0.69 | 0.70 |
| **B221F Day 7** | 0.74 | 0.95 | 0.80 | 0.83 | 0.90 | 0.93 | 0.78 | 0.78 | 0.78 | 0.68 | 0.78 | 0.82 | 0.88 | 0.74 | 0.71 | | 0.62 | 0.71 | 0.70 | 0.68 | 0.74 | 0.73 |
| **B344K Day 7** | 0.59 | 0.91 | 0.66 | 0.72 | 0.80 | 0.87 | 0.64 | 0.68 | 0.58 | 0.49 | 0.66 | 0.77 | 0.77 | 0.60 | 0.52 | 0.52 | | 0.48 | 0.53 | 0.38 | 0.56 | 0.54 |
| **D117A Day 7** | 0.79 | 1.00 | 0.85 | 0.91 | 0.96 | 0.96 | 0.85 | 0.89 | 0.78 | 0.77 | 0.85 | 0.93 | 0.96 | 0.82 | 0.78 | 0.76 | 0.61 | | 0.75 | 0.65 | 0.80 | 0.75 |
| **D414H Day 7** | 0.80 | 0.99 | 0.85 | 0.89 | 0.96 | 0.94 | 0.84 | 0.86 | 0.84 | 0.73 | 0.82 | 0.89 | 0.95 | 0.80 | 0.76 | 0.74 | 0.67 | 0.75 | | 0.72 | 0.80 | 0.80 |
| **D438H Day 7** | 0.71 | 0.94 | 0.74 | 0.80 | 0.86 | 0.91 | 0.75 | 0.77 | 0.65 | 0.61 | 0.75 | 0.84 | 0.84 | 0.67 | 0.62 | 0.63 | 0.42 | 0.57 | 0.63 | | 0.67 | 0.65 |
| **Vaccine 2017** | 0.76 | 0.98 | 0.86 | 0.94 | 1.01 | 0.98 | 0.93 | 0.91 | 0.84 | 0.78 | 0.87 | 0.94 | 1.01 | 0.91 | 0.85 | 0.82 | 0.74 | 0.83 | 0.83 | 0.79 | | 0.67 |
| **Vaccine 2018** | 0.76 | 0.96 | 0.82 | 0.91 | 0.96 | 0.96 | 0.90 | 0.86 | 0.82 | 0.78 | 0.87 | 0.92 | 0.97 | 0.86 | 0.84 | 0.78 | 0.69 | 0.76 | 0.80 | 0.74 | 0.65 | |

Fig S6. Relative L1-norm showing the genetic distance between samples. Rows give the genetic distance between a sample relative to each sample in the column.
